# miqoGraph: Fitting admixture graphs using mixed-integer quadratic optimization

**DOI:** 10.1101/801548

**Authors:** Julia Yan, Nick Patterson, Vagheesh Narasimhan

## Abstract

Admixture graphs represent the genetic relationship between a set of populations through splits, drift and admixture. In this paper we present the `Julia` package `miqoGraph`, which uses mixed-integer quadratic optimization to fit topology, drift lengths, and admixture proportions simultaneously. Inference of topology is particularly powerful, with integer optimization automating what is usually an arduous manual process.

**Availability:** https://github.com/juliayyan/PhylogeneticTrees.jl

## 1 Introduction

The genetic relationship between a set of populations cannot be described precisely by a simple tree because of the presence of admixture. An admixture graph provides a way to represent the complex relationship between populations, including their separation, subsequent drift, and possible merging by using distributions of multiple trees. Several methods exist to build and visualize admixture graphs as well as to infer optimal parameters of drift lengths and admixture edges, such as `TreeMix` (Pickrell and Pritchard, 2012), `AdmixTools` (Patterson *et al.*, 2012), `MixMapper` (Lipson *et al.*, 2013), and `admixturegraph` (Leppälä *et al.*, 2017). Some of these methods cannot simultaneously infer the optimal topology with the parameters of the graph under that topology. Rather, they require that the topology of a particular graph be pre-specified, and then infer the graph parameters. Other methods such as `TreeMix` (Pickrell and Pritchard, 2012) and `MixMapper` (Lipson *et al.*, 2013) search a restricted space of possible admixtures. In Leppälä *et al.* (2017), all possible topologies are enumerated using exhaustive searches; however such an approach becomes intractable at larger problem sizes.

Here, we describe the implementation of the `miqoGraph` package, which can simultaneously infer the optimal graph topology, drift lengths, and admixture proportions. We test our algorithm on simulated and real datasets comprising several populations and show that even as fewer parameters are specified *a priori*, running times are sped up over competitive algorithms by orders of magnitude: high-quality and optimal solutions can be found in seconds.

## 2 Methods

A typical approach to admixture graph fitting is to first specify a topology, and then compute the graph’s fit to genetic data. Drift patterns in the data can be summarized by *f*-statistics (Patterson *et al.*, 2012), and for a given topology it is possible to construct a basis set of expected values of *f*-statistics that define the graph (Pickrell and Pritchard, 2012). We will refer to the vector of empirical *f*-statistics as **f**, and given a topology **x**, drift lengths **w**, and admixture proportions ***α***, we will call the vector of the expected *f*-statistics *g*(**w**, ***α***; **x**). Drift lengths and admixture proportions are then selected to maximize the likelihood as follows:

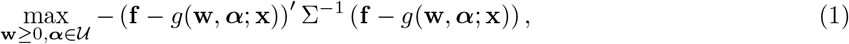

where Σ^−1^ is the covariance matrix of the empirical statistics **f**, and 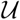 represents the set of all valid admixture proportions. This is the approach of `qpGraph`, developed by Patterson *et al.* (2012).

In our approach, which we call `miqoGraph`, rather than fixing the topology **x** before solving for drift lengths **w** and admixture proportions ***α***, we optimize over all three *simultaneously*:

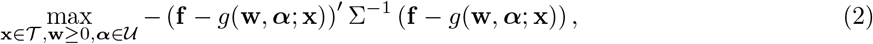

where 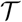 represents the set of all valid topologies. Topologies have previously been explored by enumeration over small graphs (Leppälä *et al.*, 2017), but this approach is intractable for larger graphs. Here we present a novel formulation of the problem using mixed-integer quadratic optimization (MIQO), where we model the problem of determining a best-fit graph topology as assignment of populations to leaf nodes of a binary tree. Although such problems are difficult in theory, modern optimization solvers such as Gurobi (Gurobi Optimization, Inc., 2016) and CPLEX (IBM ILOG CPLEX Optimization Studio, 2013) can quickly solve large-scale MIQO problems in practice. For an overview of integer optimization, see Wolsey and Nemhauser, 2014, and for our formulation, see the supplementary material.

Our approach requires pre-specification of the following parameters:

1. Tree depth *D* ∈ ℤ^+^,
2. Number of admixture events *A* ∈ {0} ∪ ℤ^+^, and
3. Admixture resolution *K* for *K* ∈ ℤ^+^ (only needed if *A >* 0).

If there are no admixture events (*A* = 0), the populations’ relationship can be represented using a single binary tree. We model admixture (*A >* 0) by allowing the population assignments to leaf nodes to be between 0 and 100%, and the admixture resolution *K* allows admixture proportions to be estimated to an accuracy of 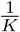. For example, *K* = 10 allows values of 0%, 10%,…, 90%, 100% (see supplementary materials).

We then solve optimization problem (2) to find the best-fit tree topology, drift lengths, and admixture proportions under the specified parameters. A major benefit of `miqoGraph` over prior approaches is the flexibility of the parameters, with each specification of parameter values representing numerous potential admixture graphs. As such, although it is computationally intractable to enumerate over the thousands of potential topologies for several populations, our algorithm quickly finds well-fit topologies using MIQO. Although it may not be obvious which parameter values are appropriate *a priori*, multiple optimization problems can be solved in parallel on a reasonable range of parameter values. In our experiments, we found that trying one tree depth, several admixture resolutions, and a few admixture events were sufficient to find the correct admixture graph topologies.

Although it is not required, prior knowledge can reduce the solution space and speed up the solution time. For example, a user can specify that the path from the root to a particular population does not contain admixture, which we found to be a particularly useful feature in our simulations.

## 3 Computational Results

In Section 3.1, we validate our models on simulated admixture graphs, before turning to real data in Section 3.2. Our methods were implemented using the Julia programming language (Bezanson *et al.*, 2014), using the optimization modeling package JuMP (Lubin and Dunning, 2015). We used the Gurobi solver version 8.0.1 (Gurobi Optimization, Inc., 2016). Computational experiments were run on a desktop computer with a 16-core Intel Xeon E5-2650 CPU, 3.40 GHz processor, and 64GB of memory. Our code is available on GitHub as the package `PhylogeneticTrees.jl`. Documentation is available in the supplementary materials.

### 3.1 Simulated Data

We tested `miqoGraph` on three simulated admixture graphs that represent the possible varieties of admixture events to validate its performance. Parameters for the simulation are described in the supplementary material.

A summary of our simulated graphs is shown in Figure 1. Figure 1a shows the base graph upon which the three simulated graphs are built. The first of these, in Figure 1b, involves a single admixture event where Population 3 is produced from equal mixtures of Populations Slot0 and Slot1. The next graph in Figure 1c is identical to the first except that the admixture proportions are changed to 10% and 90%. A more complex graph in Figure 1d has Population 3 produced from a nested admixture event between Populations Slot0, Slot1, and Slot2. These graphs will be referred to as *SimpleMix*, *UnevenMix*, and *NestedMix*, respectively. These simulations are by no means exhaustive. Because the underlying optimization model’s size scales quadratically with the admixture granularity *K* (see supplementary materials), `miqoGraph` may not be appropriate for detecting low admixture proportions; the lowest that we test is 10% in the UnevenMix case. Furthermore, our formulation models admixture only at the leaf nodes: for example, the NestedMix case is represented as Population 3 being assigned 25% to Slot0, 25% to Slot1, and 50% to Slot2. Although the nesting is straightforward in this case, `miqoGraph` may produce less interpretable results for admixture graphs with many interdependent nested admixture events. Nonetheless, `miqoGraph` is a powerful tool on a variety of use cases, as we will show.

**Figure 1:**
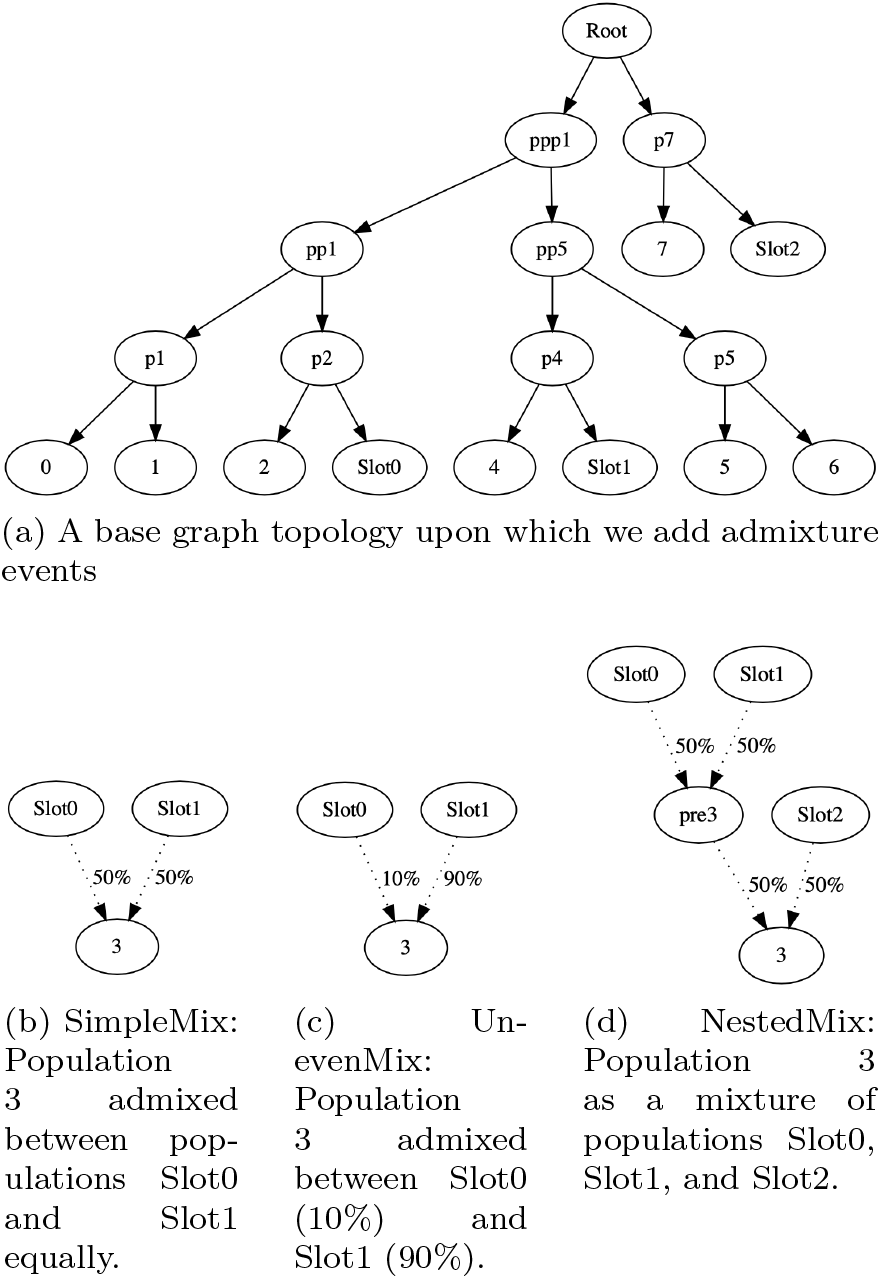
Simulated admixture graph topologies.

#### 3.1.1 The SimpleMix example

On the SimpleMix dataset, we fitted a tree with a single admixture event (*A* = 1) at an admixture resolution of *K* = 2. Our `miqoGraph` algorithm found the optimal solution in 94s with an objective value of 18.99, although it took significantly longer to prove that the topology and parameters were optimal, terminating in 1,040s. In practice, it is often the case that optimization algorithms find the optimal solution quickly but take longer to prove optimality. As such, common practice is to terminate the solver early when progress in the objective slows or halts.

However, if a guarantee of optimality is desired, it can be efficiently obtained through a combination of optimization and grid search, since the model’s solution time can speed up dramatically if it is known *a priori* that certain populations do not have admixture events in their paths to the root. We were able to perform a grid search in a total of 106s that produced the objective-18.99 tree along with a guarantee of optimality. Details of this grid search are in the supplementary material.

Most importantly, the optimal solution matched the simulated SimpleMix graph perfectly in topology and parameters.

#### 3.1.2 The UnevenMix example

In the SimpleMix case, the actual graph was admixed at exactly 50% and 50%, allowing for a low admixture granularity of *K* = 2 to capture the correct admixture event. A natural question arises when considering admixture proportions that require a higher level of resolution to capture: what level of resolution is sufficient to infer the correct admixture graph? To answer this question, we turn to the UnevenMix case of Figure 1c, where the admixture proportions are 10% and 90%.

As in the SimpleMix case, for the UnevenMix case we began with solving for a tree without admixture. In this case, `miqoGraph` terminated in 3s with an objective value of 22.08. We then ran `miqoGraph` varying the admixture resolution from *K* = 2, 3,…, 10, specifying that only Population 3 was admixed. At *K* = 7, the objective and topologies converged to the correct values and the admixture proportions reached the approximate values of 14% and 86%. This result indicates that the resolution need not be set exactly to the level required for the correct graph (*K* = 10), although it should be reasonably close. A reasonable approach might be to run `miqoGraph` at increasingly fine granularities until the topology converges. A continuous optimization algorithm such as `qpGraph` can also be run for fine-tuning.

#### 3.1.3 The NestedMix example

Our final simulated case was called NestedMix, and was similar to SimpleMix and UnevenMix except that it included a second admixture event. Admixed graphs were inferred with two admixture events, and only Population 3 was allowed to experience admixture. For both admixture resolutions *K* = 3 and 4, the inferred topology once again matched the original topology, albeit with different admixture proportions, and these trees were proved to be optimal in 33s and 63s respectively.

Table 1 shows a summary of the running times of `miqoGraph` for each dataset, compared with the running time of `qpGraph` (with the topology fixed to the correct topology). Our algorithm, `miqoGraph`, is able to find the correct graphs orders of magnitude more quickly than `qpGraph`, and with the exception of the UnevenMix case, we prove optimality more quickly as well. Our algorithm accomplishes these improved running times while also allowing exploration of varied graph topologies, when by comparison, `qpGraph` requires fixing a single graph topology *a priori*.

**Table 1:** A comparison of running times of `miqoGraph` and `qpGraph` on three simulated admixture graphs. The asterix (*) indicates where `miqoGraph` found the correct topology and parameters, but was unable to prove optimality within five minutes.

| Example | Running Time (s) |  |  |
| --- | --- | --- | --- |
|  | **miqoGraph**<br>*Correct Soln.* | **miqoGraph**<br>*Prove Opt.* | **qpGraph** |
| SimpleMix ($K = 2$) | 2 | 6 | 393 |
| UnevenMix ($K = 7$) | 42 | *300 | 392 |
| NestedMix ($K = 4$) | 8 | 63 | 407 |

The three cases SimpleMix, UnevenMix, and NestedMix are toy examples, but they represent a range of common cases. In the following section, we demonstrate the performance of `miqoGraph` on real data.

### 3.2 Real Data from Eurasia and the Americas

After testing the performance and correctness of `miqoGraph` on simulated data, we ran it on a six-population of modern and ancient DNA samples from Eurasia and the Americas to infer the phylogeny of populations leading to the Karitiana, a South American population from Brazil.

Even without specifying which population should be admixed and at a coarse admixture granularity of *K* = 2, `miqoGraph` found a solution within 8s and verified optimality after 9s. Most importantly, the graph matched one found using exhaustive searches with `qpGraph`.

We were able to refine the admixture proportions by running `miqoGraph` at higher resolutions of *K* = 3 and 4. By leveraging the knowledge that Karatiana should be admixed, learned from the *K* = 2 output, `miqoGraph` inferred the correct topology almost instantaneously (1s and 3s, respectively). At *K* = 4, the inferred admixture proportions 25%-75% corresponded closely to the values of 28%-72% estimated by `qpGraph`, and the drift lengths were also similar (see supplementary materials). The admixture graph inferred by `miqoGraph` at *K* = 4 is shown in Figure 2. In this topology, Karatiana is admixed between an ancient North Eurasian-related and a present-day East Asian-related source, consistent with previous results examining the initial peopling of the Americas (Raghavan *et al.*, 2014).

**Figure 2:**
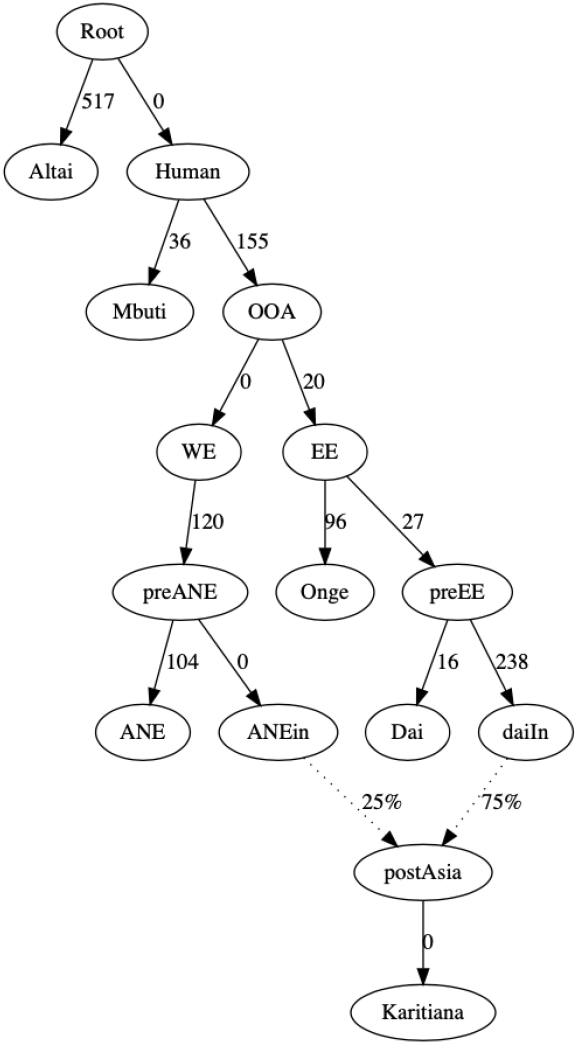
Topology, drift lengths and admixture proportions inferred on a dataset from Eurasia and the Americas.

## 4 Limitations

The main limitation of `miqoGraph` lies in the restriction of admixture events to the leaf nodes of the graph and therefore, the interpretation of its output in the presence of multiple nested admixture events. Suppose a particular population A has admixture from populations B and C, and that B itself is admixed from D and E. The ordering of these events is not captured in our representation of the graph, and it can be challenging to reconstruct the correct sequence of events leading to the true admixture graph. To aid interpretability, our framework allows the user to sequentially add new populations while fixing the topology for other populations. The positions of these new populations can vary freely, or they can be tentatively assigned to positions based on the user’s best guess, giving the optimizer a “warm start” to improve upon. A second issue with our approach is that the proportion of admixture inferred is done in discrete values whose granularity is specified *a priori*. It is possible that at low admixture granularities, the best-fit topology may be incorrect. One possible way to mitigate this effect is to use `miqoGraph` to explore a possible set of graph topologies and then to use continuous optimizers such as that implemented in `AdmixTools` (Patterson *et al.*, 2012) to fit parameters on these topologies.

## 5 Conclusion

Our results show that `miqoGraph` is able to simultaneously infer topologies, drift lengths, and admixture proportions in seconds to minutes for both simulated and real-world cases. The formulation is primarily useful in settings with few nested admixture events. Nonetheless, the use of integer optimization to model what was previously a combination of labor-intensive manual enumeration and continuous optimization represents a significant step forward in efficient inference of admixture graphs.

## Acknowledgements

The authors thank Dimitris Bertsimas and members of the Reich laboratory for productive discussions.

