## Supplementary Online Material for "miqoGraph: Fitting admixture graphs using mixed-integer quadratic optimization"

### `miqoGraph` Supplementary Material

Section 1 provides documentation on installation and running the algorithm. Section 2 reviews some preliminaries of $f$-statistics and admixture. Section 3 explains the details of the mixed-integer quadratic optimization formulation. Section 4 includes the parameters used for simulating the SimpleMix, UnevenMix, and NestedMix datasets. Section 5 provides detailed computational results.

### 1 Documentation

#### 1.1 Installation Instructions

1. Download the **Julia language** from `https://julialang.org/downloads/`. This package was developed using v1.0, but has also been tested on v1.1. Open Julia and you should see a window that looks similar to Figure 1. Documentation for Julia can be found here: `https://docs.julialang.org/en/v1/index.html`. Directions for running Julia directly from the terminal can be found here: `https://en.wikibooks.org/wiki/Introducing_Julia/Getting_started#Running_directly_from_terminal`.

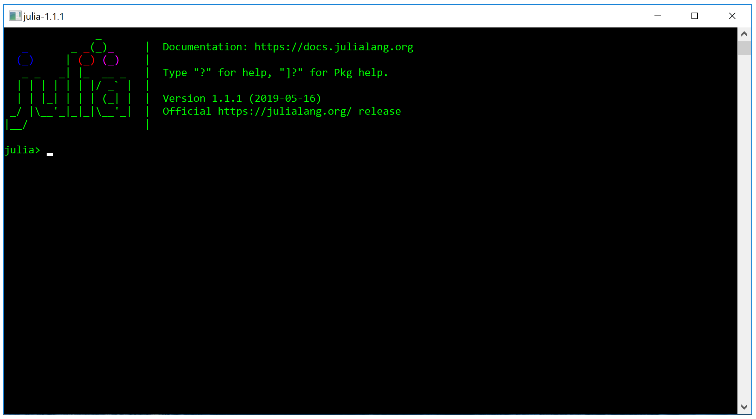


Figure 1: The Julia terminal.

2. Download the **Gurobi Optimizer**. We provide a summary of the necessary instructions here; for further issues, consult the quick start guide: `https://www.gurobi.com/documentation/quickstart.html` (**Note:** this is more comprehensive than what you will need. For example, you should not need to follow instructions for interfacing with other languages).

   (a) **Students, faculty, and staff at degree-granting academic institutions.** You must be on a university network and use your university email address for this step. Register for an academic license here: `https://www.gurobi.com/downloads/end-user-license-agreement-academic/`. Then log in. Go to the downloads page: `https://www.gurobi.com/downloads/`. Download the Gurobi Optimizer. Navigate back to the downloads page, and under **Request a License**, click Academic License. Accept the conditions, and run the `grbgetkey` command given at the bottom of the page under **Installation**. The license needs to be renewed after one year.

   (b) **Other users.** A free trial of Gurobi is available here: `https://www.gurobi.com/free-trial/`. We are working on compatibility with open-source solvers.

   If you installed Gurobi to a non-default location you need to set several environment variables. On a bash shell you can do this by adding the following lines to your .bashrc file. Once these are added, please re-login to your shell to set these or export them in your current session.

```
export GUROBI_HOME="/user/gurobi801/linux64"
export PATH="${PATH}:${GUROBI_HOME}/bin"
export LD_LIBRARY_PATH="${GUROBI_HOME}/lib"
```

export GRB_LICENSE_FILE="/user/gurobi801/linux64/bin/y/gurobi.lic"
Please change these according to the relevant locations for your installation. GUROBI_HOME should be set to the location of gurobi.sh, and GRB_LICENSE_FILE should be set to the location of gurobi.lic. You can now test your license by opening the Gurobi Interactive Shell. On Windows, this can be done by opening the Gurobi Interactive Shell application or double clicking on the desktop Gurobi icon. On a Mac or on Linux, this can be done by typing gurobi.sh in the Terminal. The shell should display text similar to the following:

```
Gurobi Interactive Shell, Version 8.1.1
Copyright (c) 2019, Gurobi Optimization, LLC
Type "help()" for help
gurobi>
```

More details are available in the quick start guide.

3. Within Julia, open the Julia package manager. This can be done by typing the right bracket (]) key in the Julia terminal shown in Figure 1. You should see you are in the package manager if the text in front of the cursor switches from julia> to (v1.0) pkg> or (v1.1) pkg>. Install miqoGraph by running the following command in the package manager:
   ```
   (v1.0) pkg> add https://github.com/juliayyan/PhylogeneticTrees.jl.git
   ```
   (Do not type the colored text; it is just meant to indicate that you should be in the package manager.) For a full reference to the Julia package manager, see https://julialang.github.io/Pkg.jl/v1/. The code for PhylogeneticTrees should be contained in /.julia/packages/PhylogeneticTrees/xx where xx is a unique hash code.

4. To get access to solver-specific parameters such as time limits and output flags, you will want to install the Gurobi package separately. It will also be helpful to install the JuMP, CSV, and DataFrames packages separately. Run the following command in the package manager:
   ```
   (v1.0) pkg> add Gurobi
   (v1.0) pkg> add JuMP@0.18.5
   (v1.0) pkg> add CSV@0.4.3
   (v1.0) pkg> add DataFrames@0.17.1
   ```
   (You can generally add packages using the package name instead of the git url. Since our package is not registered in the wider Julia package ecosystem, we use the git url for installation.)

5. Within the Julia package manager, test the package by running the following command:
   ```
   (v1.0) pkg> test PhylogeneticTrees
   ```
   This will take a few minutes and print a lot of output. The final message should say Testing PhylogeneticTrees tests passed. You can exit the package manager by pressing backspace.

6. **(For future reference)** If at a future point you need to update the package, you can run the following command:
   ```
   (v1.0) pkg> update PhylogeneticTrees
   ```

**Installation troubleshooting.** If you are unable to run test PhylogeneticTrees, you may have an issue with the package dependencies. Here are some common issues:

- **Build errors.** If you see an error message saying something like `Please run Pkg.build("X")` (where X is some package name), then within the Julia package manager, run the following:
  `(v1.0) pkg> build X`
  (Do not type the colored text; it is just meant to indicate that you should be in the package manager.) Note that there are no quotation marks around the package name X in the command executed within the package manager. Quit the Julia application and try again.

- **Older or newer package versions.** Within the Julia package manager, type the following command:
  `(v1.0) pkg> status`
  This package was developed under JuMP v0.18.5, CSV v0.4.3, and DataFrames v0.17.1. If you see other versions listed, you can switch to these versions with the following commands:
  ```
  (v1.0) pkg> add JuMP@0.18.5
  (v1.0) pkg> add CSV@0.4.3
  (v1.0) pkg> add DataFrames@0.17.1
  ```
  Note that if you are using Julia 1.2 or above, you will need JuMP v0.18.6 instead of JuMP v0.18.5.

#### 1.2 Usage

Some code and data examples are provided in the `test/` directory of the GitHub repository. Here, we give more detail on the required input data and code.

##### 1.2.1 Input Data

**$f_3$-statistics file.** The $f_3$-statistics are provided as a CSV. The CSV must follow the following header format:

```
Outgroup,A,B,f3
```

where the first column is the outgroup, the second and third columns are populations, and the fourth column is the value of the statistic $F_3(\texttt{Outgroup}; \texttt{A}, \texttt{B})$. Each row will be a different $f_3$-statistic, but it is not necessary to include both permutations of the `A` and `B` columns. All population names must be `Strings`, i.e., 7 is not a valid population name. They also should not include spaces.

The first few rows of a sample $f_3$-statistics CSV are shown below as an example. The remainder of the file can be found in `test/testdata/f3.Europe6.csv`.

```
Outgroup,A,B,f3
Mbuti,Altai,Altai,482.47
Mbuti,Altai,WEHG,34.748
```

**Covariance file.** The covariance matrix is also provided as a CSV. The CSV must have the following header format:

```
A1,B1,A2,B2,covariance
```

where the first four columns are populations, and the fifth column is the value of the covariance for $F_3(\texttt{Outgroup}; \texttt{A1}, \texttt{B1})$ and $F_3(\texttt{Outgroup}; \texttt{A2}, \texttt{B2})$. Each row will be a different element of the covariance matrix, but it is not necessary to include all permutations of the columns. All population names must match those in the $f_3$-statistics file.

The first few rows of a sample covariance CSV are shown below as an example. The remainder of the file can be found in `test/testdata/f3.Europe6-covariance.csv`.

```
A1,B1,A2,B2,covariance
Altai,Altai,Altai,Altai,4.531
Altai,Altai,Altai,WEHG,1.535
```

##### 1.2.2 A Default Wrapper

For user convenience, a default wrapper is contained in `example/default.jl`, along with a default parameters file in `example/params-miqo.csv`.

The `params-miqo.csv` file contains the following fields:

- `mean_file` The name of the file containing the $f_3$ statistics,
- `cov_file` The name of the file containing the covariance matrix,
- `output_file` The name of the file that PhylogeneticTrees should write output (the topology) to,
- `log_file` The name of the file that PhylogeneticTrees should write logging information to,
- `time_limit` A time limit (in seconds) for Gurobi,
- `warm_start` 1 if the model with admixture should warm-start the problem with a tree without admixture and 0 otherwise,
- `warm_start_time_limit` A time limit (in seconds) for the warm start model,
- `depth` Depth of the tree (a count of the edges from root to leaf),
- `granularity` Admixture granularity ($K \geq 1$),
- `admixture_events` Number of admixture events ($A \geq 0$),
- `unmixed_pops` Any populations (whose names should match those in the mean and covariance files) that should not experience admixture in the model, separated by spaces.

The default wrapper for the Americas example in our paper can be run by navigating to `example/` and running the command `julia default.jl params-miqo.csv`. The wrapper by default looks for a file called `params-miqo.csv`, so the argument can be omitted if this is the parameters file name.

##### 1.2.3 Reading Data

1. If you are still in the Julia package manager, exit the package manager by pressing backspace. You should see the `julia>` text before your cursor.

2. Within Julia, navigate to the directory containing your data files. Open the shell mode by typing the semicolon (;) key in the Julia terminal shown in Figure 1. You should see that you are in shell mode if the text in front of the cursor switches from `julia>` to `shell>`. Once in shell mode, you can use the system shell to execute system commands. Navigate to the directory that is storing your data files with the following command:
   `shell> cd YOUR_DIRECTORY`

3. Exit shell mode by pressing backspace. You should see the `julia>` text before your cursor. Load the `PhylogeneticTrees` and `Gurobi` packages with the following line of code:
```
julia> using PhylogeneticTrees, Gurobi
```

4. You can then read data from your files using the following code:
```
julia> pd = PhylogeneticTrees.PopulationData("F3_FILE.csv", "COVARIANCE_FILE.csv")
```
The first argument of the `PhylogeneticTrees.PopulationData()` function is the file name of the $f_3$-statistics CSV file, and the second argument is the file name of the covariance CSV file. This code stores the population data in a data structure named `pd`.

5. The number of populations, outgroup key, population names, $f_3$-statistics, and covariance matrix can be accessed through `pd.npop`, `pd.outgroup`, `pd.pops`, `pd.f3`, and `pd.cov`, respectively.

##### 1.2.4 Fitting a Model

Following these steps will allow you to construct a model **without admixture**.

1. Build a binary tree data structure of depth $D = 3$ with the following code:
```
julia> D = 3
julia> bt = PhylogeneticTrees.BinaryTree(D)
```
This will create a binary tree with $2^{D+1} - 1 = 15$ nodes, $2^D = 8$ of which are leaf nodes.

2. Construct the optimization model with the following code:
```
julia> tp = PhylogeneticTrees.TreeProblem(pd, bt, solver = GurobiSolver(TimeLimit = 60))
```
The first argument of `PhylogeneticTrees.TreeProblem()` is the data structure containing the population data, the second argument is the binary tree, and the third is the solver. The `TimeLimit` flag indicates that the solver will terminate after 60 seconds.

3. To solve the model, use the following code:
```
julia> solve(tp.model)
```
The `solve()` function comes from the JuMP package, which you should have installed separately.

4. To print which populations were assigned to which nodes, you can use the following code:
```
julia> PhylogeneticTrees.printnodes(tp)
```
In this output, the first column is the population name, the second column is the node, and the third column is the proportion of the population that was assigned to that node (always 1.0 if there is no admixture).
For relatively small trees (depth 4 and below), you can print a visualization of the tree itself using the following code:
```
julia> PhylogeneticTrees.printtree(tp)
```

Admixture events can be added using optional arguments to the `PhylogeneticTrees.TreeProblem()` function:

- `nlevels` (default `1`): the level of granularity $K$ that is desired for admixture proportion estimation. The higher $K$ is, the more precise the estimation, which is in intervals of $\frac{1}{K}$. By default, the granularity level is set to 1, indicating no admixture. We recommend starting with a low level before moving to higher levels to test what your computer can handle. We have tested up to $K = 10$, and do not recommend going significantly further. At $K = 10$, the proportions can take on values $0\%, 10\%, 20\%, \ldots, 90\%, 100\%$.

- `nmixtures` (default `pd.npop`): the maximum number of nodes that can be assigned to, or `pd.npop`$+A$, where $A$ is the number of admixture events. By default, `nmixtures = pd.npop` indicates $A = 0$, meaning that there are no admixture events.

Now, to construct a model **with admixture**, we model the previous procedure as follows:

1. Same as without admixture

2. Construct the optimization model with the following code:
```
julia> tp = PhylogeneticTrees.TreeProblem(pd, bt, solver = GurobiSolver(TimeLimit = 60),
nlevels = 2, nmixtures = pd.npop + 1)
```
The extra parameters allow one of the populations to be mixed at 50%-50%.

3. The mode can be solved directly as before, but a couple of extra lines of code can help the model solve more quickly.

   (a) A "warm start" can be provided to the solver using the following code:
   ```
   julia> PhylogeneticTrees.warmstartunmixed(tp, timelimit=30)
   ```
   This finds the best possible tree without admixture within the time limit specified by the (optional) `timelimit` parameter (in seconds, with a default of 30 seconds). Then it loads the tree without admixture as a starting solution before attempting to add admixture.

   (b) Populations that should not be admixed can be specified using the following code:
   ```
   julia> PhylogeneticTrees.unmix(tp, "POPULATION")
   ```
   where the name of the population is provided as a `String` in the second argument to `PhylogeneticTrees.unmix()`. This name must match what is in `pd.pops`.

4. Same as without admixture

**Model-fitting troubleshooting.** Here are some common issues:

- **Data input.** If you see an error message saying something like `Objective Q not PSD`, then you have probably inputted your covariance matrix incorrectly. Check the formatting instructions and try again.

##### 1.2.5 Running Scripts

If you have saved all your previous commands in a script called `script.jl`, you can save yourself some typing by using the following code to execute the commands of the script:
```
julia> include("script.jl")
```

### 2 Preliminaries

#### 2.1 $f$-Statistics

Let $\mathcal{P}$ represent a set of populations. Let $X_p$ denote the random variable corresponding to the allele frequencies at a single polymorphism in population $p \in \mathcal{P}$. We also have a tree $\mathcal{T}$ that is composed of nodes $\mathcal{V}$ and edges $\mathcal{E}$. In general we use the indices $p, q, r, s, u, v$ to refer to populations, and $i, j$ to refer to general nodes.

Allele frequencies are considered to follow a *martingale property*. In particular, if the edge $(p, q)$ is present in the graph, meaning that $q$ is a descendant of $p$, then we have

$$\mathbb{E}[X_q | X_p = x] = x. \tag{1}$$

The $f_2$-statistic, also called *branch length*, is the squared drift. For two populations $p$ and $q$, it is defined as follows:

$$F_2(p, q) = \mathbb{E}[(X_p - X_q)^2]. \tag{2}$$

It is assumed that drifts on distinct edges of the phylogenetic tree are orthogonal. Namely, for two distinct edges $(p, q)$ and $(r, s)$, we have

$$\mathbb{E}[(X_p - X_q)(X_r - X_s)] = 0. \tag{3}$$

This property means that $f_2$-statistics (branch lengths) are additive. Consider a path $p \to q \to r$ in our tree. We can show the additivity of branch lengths as follows:

$$\begin{aligned} F_2(r, p) &= \mathbb{E}[(X_r - X_p)^2] \\ &= \mathbb{E}[(X_r - X_q + X_q - X_p)^2] \\ &= \mathbb{E}[(X_r - X_q)^2] + 2\mathbb{E}[(X_r - X_q)(X_q - X_p)] + \mathbb{E}[(X_q - X_p)^2] \\ &= \mathbb{E}[(X_r - X_q)^2] + \mathbb{E}[(X_q - X_p)^2] \\ &= F_2(r, q) + F_2(q, p). \end{aligned}$$

The rest can be shown by induction.

The $f_3$-statistic, for three populations $p$,$q$, and $r$, is defined as follows:

$$F_3(p; q, r) = \mathbb{E}[(X_p - X_q)(X_p - X_r)]. \tag{4}$$

The $f_3$-statistics can also be computed as sums of the $f_2$ statistics by inspecting the paths from population $r$ to population $p$ in the tree, and similarly for population $q$ to population $p$ (ignoring edge direction).

$$\begin{aligned} F_3(p; q, r) &= \mathbb{E}[(X_p - X_q)(X_p - X_r)] \\ &= \mathbb{E}\left[\left(\sum_{(u,v)\in\mathcal{E}} (X_u - X_v)\mathbb{1}_{\{(u,v)\in \mathrm{path}(q,\,p)\}}\right)\left(\sum_{(u,v)\in\mathcal{E}} (X_u - X_v)\mathbb{1}_{\{(u,v)\in \mathrm{path}(r,\,p)\}}\right)\right] \end{aligned}$$

$$
\begin{aligned}
&= \mathbb{E}\left[\left(\sum_{(u,v)\in\mathcal{E}} (X_u - X_v)^2 \mathbb{1}_{\{(u,v)\in \mathrm{path}(q,p)\cap \mathrm{path}(r,p)\}}\right)\right] \\
&= \sum_{(u,v)\in\mathcal{E}} \mathbb{E}\left[(X_u - X_v)^2\right] \mathbb{1}_{\{(u,v)\in \mathrm{path}(q,\,p)\cap \mathrm{path}(r,\,p)\}} \\
&= \sum_{(u,v)\in\mathcal{E}} F_2(u,v) \mathbb{1}_{\{(u,v)\in \mathrm{path}(q,\,p)\cap \mathrm{path}(r,\,p)\}}, \qquad (5)
\end{aligned}
$$

where $\mathrm{path}(q,p)$ indicates the sequence of edges on the path from the node containing population $q$ to the node containing population $p$, ignoring edge direction. Namely, the statistic $F_3(p;q,r)$ is the sum of the branch lengths of the overlapping edges of the paths from $q$ to $p$ and $r$ to $p$. Note that $F_3(p;q,q) = F_2(p,q)$.

Typically, $f_3$-statistics are collected with an *outgroup* as the first argument. An outgroup is a population that split up from the ancestral populations of the other populations of study, before the population splits leading to the other populations.

#### 2.2 Admixture

If two populations $p$ and $q$ are admixed with mixing proportions $\alpha$ and $1-\alpha$, respectively, then the resulting allele frequency will be

$$
\alpha X_p + (1-\alpha) X_q. \qquad (6)
$$

Given the mixing relationship (6), we can derive the theoretical formulas for admixed populations. For example, if population $p$ is mixed from populations $q$ and $r$ at frequencies $\alpha$ and $(1-\alpha)$ respectively, and both $q$ and $r$ do not experience admixture, then $F_2(o,p)$ (with outgroup $o$) can be calculated as follows:

$$
\begin{aligned}
F_2(o,p) &= \mathbb{E}[(\alpha X_q + (1-\alpha)X_r - X_o)^2] \\
&= \mathbb{E}[(\alpha(X_q - X_o) + (1-\alpha)(X_r - X_o))^2] \\
&= \alpha^2 \mathbb{E}[(X_q - X_o)^2] + 2\alpha(1-\alpha)\mathbb{E}[(X_q - X_o)(X_r - X_o)] + (1-\alpha)^2 \mathbb{E}[(X_r - X_o)^2] \\
&= \alpha^2 F_2(o,q) + 2\alpha(1-\alpha) F_3(o;q,r) + (1-\alpha)^2 F_2(o,r) \\
&= \alpha^2 \sum_{(u',v')\in \mathrm{path}(o,q)} F_2(u',v') \\
&\quad + 2\alpha(1-\alpha) \sum_{(u',v')\in \mathrm{path}(o,q)\cap \mathrm{path}(o,r)} F_2(u',v') \\
&\quad + (1-\alpha)^2 \sum_{(u',v')\in \mathrm{path}(o,r)} F_2(u',v'). \qquad (7)
\end{aligned}
$$

We now provide a general statement on $f_3$-statistics with admixture. For brevity, we will refer to populations that do not experience admixture in their path to the root as "unmixed." If population $p$ is mixed from unmixed ancestral populations $r \in \mathcal{A}_p$ at levels $\{\alpha_r\}_{r\in\mathcal{A}_p}$ ($\alpha_r \geq 0$ and $\sum_r \alpha_r = 1$) and population $q$ is mixed from unmixed ancestral populations $s \in \mathcal{A}_q$ at levels $\{\beta_s\}_{s\in\mathcal{A}_q}$ ($\beta_s \geq 0$ and $\sum_s \beta_s = 1$), the statistic $F_3(o;p,q)$ can be computed as follows:

$$
F_3(o;p,q) = \mathbb{E}\left[\left(\sum_{r\in\mathcal{A}_p} \alpha_r X_r - X_o\right)\left(\sum_{s\in\mathcal{A}_s} \beta_s X_s - X_o\right)\right]
$$

$$
\begin{aligned}
&= \mathbb{E}\left[\left(\sum_{r\in\mathcal{A}_p}\alpha_r(X_r - X_o)\right)\left(\sum_{s\in\mathcal{A}_s}\beta_s(X_s - X_o)\right)\right] \\
&= \sum_{r\in\mathcal{A}_p}\sum_{s\in\mathcal{A}_s}\alpha_r\beta_s\mathbb{E}\left[(X_r - X_o)(X_s - X_o)\right] \\
&= \sum_{r\in\mathcal{A}_p}\sum_{s\in\mathcal{A}_s}\alpha_r\beta_s\left(\sum_{(u,v)\in\text{path}(r,o)\cap\text{path}(s,o)} F_2(u,v)\right). \qquad (8)
\end{aligned}
$$

Note equation (8) can also represent nested layers of two-way admixtures, since any admixture topology can be represented as a mixture of multiple unmixed ancestral populations. For example, if population $q$ is 50% population $p$ and 50% population $r$, and population $r$ is in turn 50% population $s$ and 50% population $u$, then population $q$ is 50% population $p$, 25% population $s$, and 25 population $u$.

In our formulation, it will be more convenient to rearrange the terms of (8). An edge $(u, v)$ is in $\text{path}(r, o) \cap \text{path}(s, o)$ if it satisfies certain conditions. There are two cases. First, if $(u, v)$ *is not* on the path from the root to the outgroup, then populations $r$ and $s$ must be descendants of $v$, or $v$ itself. Second, if $(u, v)$ *is* on the path from the root to the outgroup, then populations $r$ and $s$ cannot be descendants of $v$, or $v$ itself. Therefore, equation (8) is equivalent to the following:

$$
\begin{aligned}
F_3(o; p, q) = &\sum_{(u,v)\in\mathcal{E}\setminus\text{path}(\text{root},o)} F_2(u,v) \sum_{r\in\mathcal{A}_p\cap\mathcal{D}_v}\sum_{s\in\mathcal{A}_q\cap\mathcal{D}_v}\alpha_r\beta_s + \\
&\sum_{(u,v)\in\text{path}(\text{root},o)} F_2(u,v) \sum_{r\in\mathcal{A}_p\setminus\mathcal{D}_v}\sum_{s\in\mathcal{A}_q\setminus\mathcal{D}_v}\alpha_r\beta_s. \qquad (9)
\end{aligned}
$$

### 3 Integer Optimization Formulation

We can now formulate a mixed-integer quadratic optimization problem to solve for an optimal tree. We first study the case without admixture in Section 3.1, before generalizing to admixed populations in Section 3.2.

#### 3.1 Without Admixture

We start with the following **input data**:

- $\mathcal{P}$: A set of populations
- $o \in \mathcal{P}$: An outgroup
- $\hat{f}_{p,q}$: $f_3$-statistics $F_3(o;p,q)$ measured relative to the outgroup.

We also specify the following tree parameters:

- $D \in \mathbb{Z}_+$: Desired depth of the tree (which will have nodes $1, 2, \ldots, 2^{D+1} - 1$)
- $n = 2^D$: The node at which to fix the outgroup.

The depth of the tree should be chosen large enough so that the populations $\mathcal{P}$ can comfortably fit in the $2^D$ leaf nodes. The choice of $n = 2^D$ is arbitrary; any leaf node $2^D, 2^D + 1, \ldots, 2^{D+1} - 1$ would suffice.

Our optimization problem will have the following **main decision variables** that parametrize the tree:

- $x_{p,i} \in \{0, 1\}$: 1 if population $p$ is assigned to node $i$, 0 otherwise. We only include nodes $i$ that are leaves of the tree, which we will refer to as $\mathcal{V}_D := \{2^D, \ldots, 2^{D+1} - 1\}$. The root node is labeled 1.
- $w_{i,j} \geq 0$: branch length on edge $(i, j)$. This is also the $f_2$-statistic of equation (5).

In order to fit the $f$-statistics, the following **auxiliary decision variables** will also be needed:

- $y_{p,q,(i,j)} \geq 0$: although not explicitly defined to be binary, will be constrained to take on the value 1 if edge $(i, j)$ contributes to the $F_3(o;p,q)$ statistic, and 0 otherwise.
- $z_{p,q,(i,j)} \geq 0$: the branch length on edge $(i, j)$ if it contributes to $F_3(o;p,q)$, and 0 otherwise.

Finally, the $f$-statistics and errors will be computed using the following **fitting decision variables**:

- $f_{p,q} \geq 0$: the fitted value of $F_3(o;p,q)$
- $\epsilon_{p,q}$: the error associated with fitting $F_3(o;p,q)$.

We now turn to the constraints necessary to model the problem. First, the outgroup must be assigned to the proper node:

$$x_{o,n} = 1. \tag{10}$$

Each population is assigned to a leaf node using the following constraint:

$$\sum_{i \in \mathcal{V}_D} x_{p,i} = 1 \quad \forall p \in \mathcal{P}. \tag{11}$$

Similarly, each leaf node is assigned at most one population as follows:

$$\sum_{p \in \mathcal{P}} x_{p,i} \leq 1 \quad \forall i \in \mathcal{V}_D. \tag{12}$$

Using $\mathcal{S}_j$ to denote the leaves of the subtree rooted at $j$ and recalling equation (9) and the definition of $\mathbf{y}$, we use the following constraints to link the auxiliary $\mathbf{y}$ and main $\mathbf{x}$ variables:

$$y_{p,q,(i,j)} \leq \sum_{j' \in \mathcal{S}_j} x_{p,j'} \tag{13a}$$

$$y_{p,q,(i,j)} \leq \sum_{j' \in \mathcal{S}_j} x_{q,j'} \tag{13b}$$

$$y_{p,q,(i,j)} \geq \sum_{j' \in \mathcal{S}_j} (x_{p,j'} + x_{q,j'}) - 1 \qquad \forall p \in \mathcal{P}, q \in \mathcal{P}, (i,j) \in \mathcal{E} \setminus \text{path}(n,1), \tag{13c}$$

for edges not in the path from the outgroup node to the root, and

$$y_{p,q,(i,j)} \leq 1 - \sum_{j' \in \mathcal{S}_j} x_{p,j'} \tag{14a}$$

$$y_{p,q,(i,j)} \leq 1 - \sum_{j' \in \mathcal{S}_j} x_{q,j'} \tag{14b}$$

$$y_{p,q,(i,j)} \geq 1 - \sum_{j' \in \mathcal{S}_j} (x_{p,j'} + x_{q,j'}) \qquad \forall p \in \mathcal{P}, q \in \mathcal{P}, (i,j) \in \text{path}(n,1). \tag{14c}$$

for edges in the path from the outgroup node to the root. These constraints are exact even without explicitly enforcing that the $\mathbf{y}$ variables be binary; if the $\mathbf{y}$ variables are constrained to be nonnegative then they will naturally only take 0-1 values.

With the $\mathbf{y}$ variables set to the appropriate values, the theoretical $F_3(o; p, q)$ statistic can be computed as follows:

$$f_{p,q} = \sum_{(i,j) \in \mathcal{E}} w_{i,j} y_{p,q,(i,j)} \qquad \forall p \in \mathcal{P}, q \in \mathcal{P}, \tag{15}$$

which is bilinear. However, because the $\mathbf{y}$ variables only take 0-1 values, we can set auxiliary variables $z_{p,q,(i,j)}$ to take on the values of $w_{i,j} y_{p,q,(i,j)}$ using the McCormick relaxation:

$$z_{p,q,(i,j)} \leq M y_{p,q,(i,j)} \tag{16a}$$

$$z_{p,q,(i,j)} \leq w_{i,j} \tag{16b}$$

$$z_{p,q,(i,j)} \geq w_{i,j} + M y_{p,q,(i,j)} - M \qquad \forall p \in \mathcal{P}, q \in \mathcal{P}, (u,v) \in \mathcal{E}, \tag{16c}$$

where $M$ is an upper bound representing the largest possible edge weight. We choose $M = \Gamma \max_{p,q} f_{p,q}$, with $\Gamma \geq 1$. Then, we can replace constraint (15) with the linear constraint

$$f_{p,q} = \sum_{(i,j) \in \mathcal{E}} z_{p,q,(i,j)} \qquad \forall p \in \mathcal{P}, q \in \mathcal{P}. \tag{17}$$

Finally, we can set the error terms using the following constraint:

$$\epsilon_{p,q} = \hat{f}_{p,q} - f_{p,q} \tag{18}$$

We seek to minimize the error (weighted by the inverse covariance matrix):

$$\min_{\mathbf{x},\mathbf{y},\mathbf{w},\mathbf{g},\boldsymbol{\epsilon}} \quad \sum_{p\in\mathcal{P},q\in\mathcal{P}} \sum_{p'\in\mathcal{P},q'\in\mathcal{P}} \Sigma^{-1}_{p,q,p',q'}\epsilon_{p,q}\epsilon_{p',q'}, \tag{19}$$

where $\Sigma^{-1}_{p,q,p',q'}$ is the entry in the inverted covariance matrix corresponding to $F_3(o;p,q)$ and $F_3(o;p',q')$.

#### 3.2 With Admixture

Admixture could be represented by relaxing the $\mathbf{x}$ variables to be continuous. (Then, it would also be desirable to constrain the $\mathbf{x}$ variables so that still only one population can be assigned to each node, which can be done using auxiliary binary variables.) However, this would mean that the $\mathbf{y}$ variables could take on continuous values in $[0,1]$, and therefore linearization that produced constraints (13) and (14) would no longer be exact. We have seen computationally that the big-$M$ bounds are not tight enough in the linear relaxation to produce reliable results.

Rather than relax the $\mathbf{x}$ variables to be continuous, we can approximate the continuous admixture proportions by choosing a suitably large $K \in \mathbb{Z}_+$ and introduce the following sum of auxiliary binary variables $\boldsymbol{\chi} \in \{0,1\}^{|\mathcal{P}|\times 2^D \times K}$:

$$\text{admixure proportion } (p,i) = \frac{1}{K}\sum_{k=1}^{K} \chi^k_{p,i}. \tag{20}$$

The summation in equation (20) represents the proportion of admixture for population $p$ arising from the ancestral population in node $i$. Of course, if the value is 1, then population $p$ is fully assigned to node $i$. If $K = 10$, then the admixture proportions can take on values $0\%, 10\%, \ldots, 90\%, 100\%$.

Constraints (10) and (11) need to be modified to accommodate these new $\boldsymbol{\chi}$ variables as follows:

$$\chi^k_{o,n} = 1 \quad \forall k = 1,\ldots,K, \tag{21}$$

$$\sum_{i\in\mathcal{V}_D} \chi^k_{p,i} = 1 \quad \forall p \in \mathcal{P},\ k = 1,\ldots,K. \tag{22}$$

Constraint (12) is still valid, but the $\mathbf{x}$ variables and $\boldsymbol{\chi}$ variables need to be linked in order to ensure that each node is assigned at most one population as follows:

$$x_{p,i} \geq \chi^k_{p,i} \quad \forall p \in \mathcal{P},\ i \in \mathcal{V}_D,\ k \in 1,\ldots,K, \tag{23}$$

which ensures that if $\chi^k_{p,i} = 1$ for any $k = 1,\ldots,K$, then $x_{p,i} = 1$ as well. Additionally, if we want to restrict the number of admixture events to be at most some number $A$, we can add the following constraint:

$$\sum_{p\in\mathcal{P}} \sum_{i\in\mathcal{V}_D} x_{p,i} \leq |\mathcal{P}| + A. \tag{24}$$

Because the new variables $\boldsymbol{\chi}$ are binary, we can replace the auxiliary variables $\mathbf{y}$ with auxiliary variables

$\boldsymbol{\psi} \in [0,1]^{|\mathcal{P}|^2 \times |\mathcal{E} \times K^2}$, where each variable $\psi^{k,\ell}_{p,q,(i,j)}$ is related to $\chi^k_{p,j'}$ and $\chi^\ell_{q,j'}$ for $j' \in \mathcal{S}_j$ in the same way that $y_{p,q,(i,j)}$ was related to $x_{p,j'}$ and $x_{q,j'}$ for $j' \in \mathcal{S}_j$ (cf. constraints (13) and (14)). As before, since the $\boldsymbol{\chi}$ variables are binary, the $\boldsymbol{\psi}$ variables will also take only 0-1 values despite not explicitly being constrained to be binary. Then, substituting the expanded variables into constraint (15), we obtain the following:

$$f_{p,q} = \frac{1}{K^2} \sum_{(i,j)\in\mathcal{E}} w_{i,j} \sum_{k=1}^{K} \sum_{\ell=1}^{K} \psi^{k,\ell}_{p,q,(i,j)}, \tag{25}$$

which is the $K$-level modification of constraint (15). As before, constraint (25) can also be linearized again with a McCormick relaxation, since the $\boldsymbol{\psi}$ variables only take 0-1 values. The constraint (18) and objective (19) can remain the same.

This model has $O(|\mathcal{P}|^2 2^D K^2)$ auxiliary variables, and in the case where $K = 1$, it reduces to the formulation without admixture.

### 4 Simulated Data

We used the msprime coalescent simulator [1] to simulate chromosomes (excluding the autosomes) at their relevant lengths from a set of eight populations with different topologies. We used a mutation rate of $1.5 \times 10^{-8}$, a recombination rate of $1 \times 10^{-8}$, and Ne of 500, and sampled 20 individuals from each population. To reflect the analysis performed with real data, we then created 40 haploid chromosomes from these 20 diploid sequences. We then computed empirical $f$-statistics and their accompanying covariances using a weighted block jackknife [2].

### 5 Computational Results

#### 5.1 The SimpleMix example

The first step in recovering the SimpleMix graph was to infer a topology without admixture. The optimal topology was found in merely four seconds, with an objective value of 274.41. The topology without admixture was quite close to the actual topology, with the sole error that population 3 was placed in its ancestral pre32 node due to the graph being unable to capture admixture.

We then fitted a tree with a single admixture event at an admixture resolution of $K = 2$. Our `miqoGraph` algorithm found the optimal solution in 94 seconds with an objective value of 18.99, although it took significantly longer to prove optimality, terminating in 1,040 seconds. The optimization progression is shown in Figure 2, with time on the x-axis on a log scale, and the objective value on the y-axis. The upper bound (solid line) represents incumbent optimization solutions, with each decrease indicating that a better solution has been found. The lower bound (dotted line) represents the solver's progress towards verifying whether a solution is optimal. A solution is proved to be optimal when the upper bound meets the lower bound. Figure 2 shows that the optimization solver makes quick progress towards finding the optimal solution, but takes much longer to prove optimality. As such, common practice is to terminate the solver early.

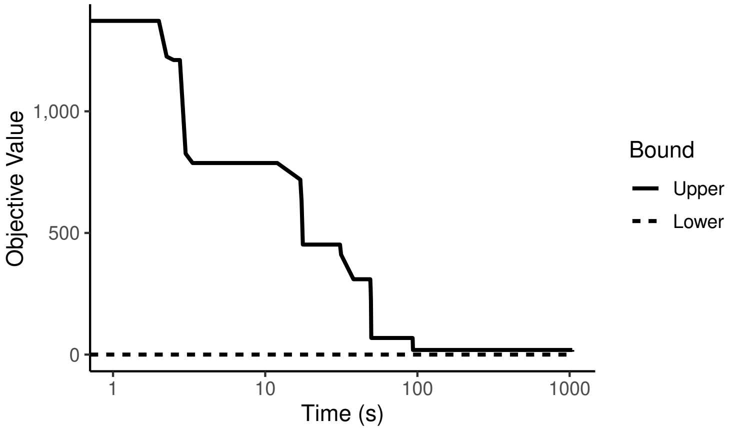


Figure 2: Optimization upper and lower bounds on the SimpleMix data with a single admixture event at a granularity of $K = 2$. Time is shown on a log scale.

If a guarantee of optimality is desired, we now demonstrate that it can be efficiently obtained through a combination of optimization and grid search. Solution time can speed up dramatically if certain populations are fixed to be unmixed *a priori*. Since we are solving for a tree with a single admixture event in the SimpleMix case, we can fix all populations to be unmixed except for one, and solve the resulting optimization problem to optimality. If we repeat this process for every population, then we can simply choose the tree with the lowest objective value, which will be optimal for a single admixture event.

The results for this grid search are shown in Table 1, and mixing population 3 clearly gives the best result that matches the objective of 18.99, verifying the optimality of the solution in the earlier model that did not have prior knowledge of which population was admixed. This enumeration takes 106 seconds total, a dramatic reduction of the running time of the original proof of optimality. The running time of this procedure can be cut further if prior knowledge is taken into account to reduce the scenarios of the grid search, since it is often the case that there are a limited number of candidates for admixture.

Table 1: Objective values and solution times on the SimpleMix data, where a single population was allowed to be admixed and all other populations were constrained to be unmixed.

| **Admixed Pop.** | **Objective** | **Time (s)** |
|---:|---:|---:|
| 1 | 274.03 | 14 |
| 2 | 274.03 | 21 |
| 3 | 18.99 | 6 |
| 4 | 274.29 | 23 |
| 5 | 67.08 | 13 |
| 6 | 274.29 | 18 |
| 8 | 248.73 | 11 |

Most importantly, the optimal solution matched the simulated SimpleMix graph perfectly in topology, illustrating the power of `miqoGraph` to quickly recover admixture graphs by optimizing topologies, weights, and mixing proportions jointly.

#### 5.2 The UnevenMix example

In the SimpleMix case, we were fortunate that the actual graph was admixed at exactly 50% and 50%, allowing for a low admixture granularity of $K = 2$ to capture the correct admixture event. A natural question arises when considering unequal admixture proportions that require a higher level of resolution to capture: what level of resolution is sufficient to infer the correct admixture graph? To answer this question, we turn to the UnevenMix case.

As in the SimpleMix case, for the UnevenMix case we began with solving for a tree without admixture. In this case, `miqoGraph` terminated in three seconds with an objective value of 22.08. We then ran `miqoGraph` varying the admixture resolution from $K = 2, 3, \ldots, 10$, specifying that only population 3 was admixed. The solution objective values for each level of resolution are shown in Figure 3 (solid line). For reference, the objective values for the correct admixed topology with the closest possible admixture weights to 10% and 90% allowed by the admixture granularity are shown as well (dotted line). For admixture resolutions $K \geq 2$, population 3 was forced to be admixed. For example, for the correct mixing proportions of 10% and 90%, the closest possible mixing proportions for a tree with granularity $K = 2$ were 50% and 50%, and for a mixed tree with granularity $K = 3$ they would be 33% and 66%. For the tree without admixture, we simply assigned population 3 to the pre32 node, which was optimal for the problem without admixture.

Because the tree without admixture at $K = 1$ is actually close in topology to the correct tree, its objective value is relatively low. However, at $K = 2$, the estimated and correct topologies differ sharply, as the 50%-50% mixing dictated by a $K = 2$ resolution is quite far from the 10%-90% reality, and as such `miqoGraph` finds another topology that differs from the correct topology but has lower objective value. Ultimately, past $K \geq 7$, the objectives and topologies converge to the correct values. This trajectory indicates that to get the right topology, the resolution need not be set exactly to the level required for the correct graph ($K = 10$), although it should be reasonably close. A reasonable approach might be to run the algorithm at increasing levels of admixture granularities until convergence in the topology is seen. A continuous optimization algorithm such as `qpGraph` can also be run to fine-tune the admixture proportions and weights.

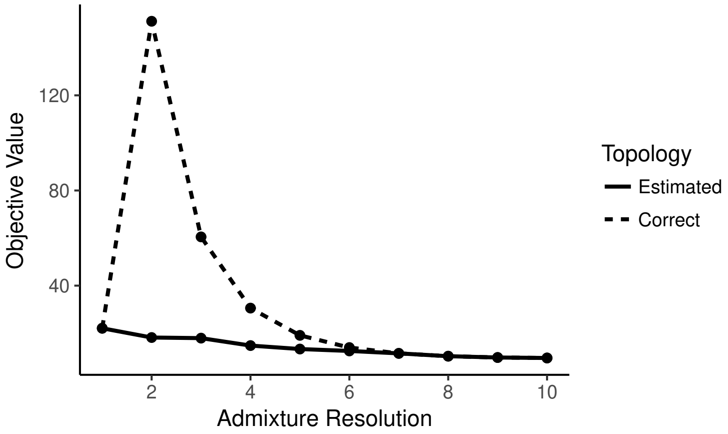


Figure 3: Objective values for increasing levels of admixture resolution on the UnevenMix case.

Formulation size and running times for varying levels of $K$ are shown in Table 2. As $K$ increases, the size of the formulation, as measured by the number of decision variables, increases dramatically. However, part of Gurobi's power is its ability to "presolve" and efficiently identify variables that can be eliminated from the model. As such, the presolved model size grows at a much more reasonable rate.

Past $K \geq 6$, Gurobi is unable to prove optimality within five minutes. However, the solutions are most likely optimal, given that the topology is close to the correct solution at $K = 6$ and is exactly correct for $K \geq 7$. The time required to find the optimal solution generally increases with $K$, as expected. However, even for $K = 10$, the solution is found in under three minutes.

Table 2: Model sizes and solution times on the UnevenMix data for varying levels of admixture granularity $K$. The asterix (*) indicates where `miqoGraph` found the correct topology and parameters, but was unable to prove optimality within five minutes.

| $K$ | **Num. Variables** | | **Running Time (s)** | |
|---|---|---|---|---|
| | *Initial* | *Presolved* | *Last Soln.* | *Prove Opt.* |
| 2 | 9,090 | 1,337 | 5 | 7 |
| 3 | 20,018 | 2,734 | 4 | 40 |
| 4 | 35,266 | 4,053 | 6 | 61 |
| 5 | 54,834 | 5,667 | 20 | 111 |
| 6 | 78,722 | 7,627 | 128 | *300 |
| 7 | 106,930 | 9,951 | 41 | *300 |
| 8 | 139,458 | 12,561 | 78 | *300 |
| 9 | 176,306 | 15,493 | 68 | *300 |
| 10 | 217,474 | 16,371 | 169 | *300 |

#### 5.3 The NestedMix example

Our final simulated case, NestedMix, followed a similar procedure as SimpleMix and UnevenMix, but had an additional complexity: in order to capture this graph, two admixture events were needed. The optimal graph without admixture was found in four seconds with an objective value of 157.16. Admixed graphs were inferred with two admixture events and all populations except for population 3 were set to be unmixed. For both admixture resolutions $K = 3$ and $K = 4$, the inferred topology once again matched the original topology exactly, albeit with different admixture proportions, and these trees were found in 33 and 63 seconds, respectively. The optimization progress for the $K = 4$ granularity is shown in Figure 4, with time shown on a log scale as in Figure 2. Even for this larger model, with the addition of the constraints restricting that only population 3 be admixed, the optimal solution is found within ten seconds, and the remainder of the time is spent proving optimality.

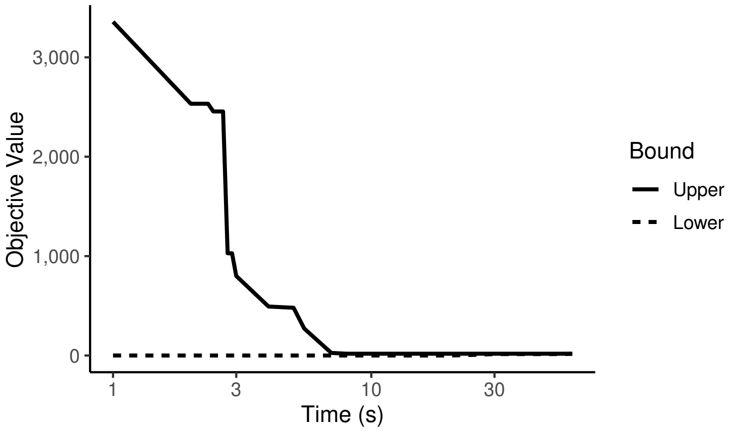


Figure 4: Optimization upper and lower bounds on the NestedMix data with two admixture events at a granularity of $K = 4$. Time is shown on a log scale.

#### 5.4 The Eurasian-American example

We ran `miqoGraph` on a six-population dataset from Eurasia and the Americas to infer the phylogeny of populations leading to the Karitiana, a South American population from Brazil. An admixture graph created using `qpGraph`, with the topology pre-specified through manual enumeration, is shown in Figure 5. Karitiana was admixed between an ancient North Eurasian-related and a present-day East Asian-related source, which is consistent with [3].

The admixture graph created using `miqoGraph` at $K = 4$ is shown in Figure 6a. Notably, the topology inferred by `miqoGraph` matches the pre-specified topology in Figure 5 (and therefore [3]), with some permutation that we explain stepwise in Figure 6. Permutation is allowed by `miqoGraph` because the $f_3$-statistics do not depend on the direction of the edges. For ease of comparison, in Figures 6b and 6c we show the steps taken to translate the output of Figure 6a into the standard format of Figure 5. The translation maps the non-leaf nodes $1-7$ of Figure 6a to an empty node, OOA, EE, Human, preANE, preEE, and an empty node, respectively. Root and WE from Figure 5 do not appear in Figure 6a, but by examining the drift lengths, we see that they were absorbed into the (4,Altai-1) and (2,5) edges, respectively. The drift of 517 along the edge (4,Altai-1) corresponds to the (Root,Altai) and (Root,Human) drifts of 258 each $(258 + 258 = 516)$.

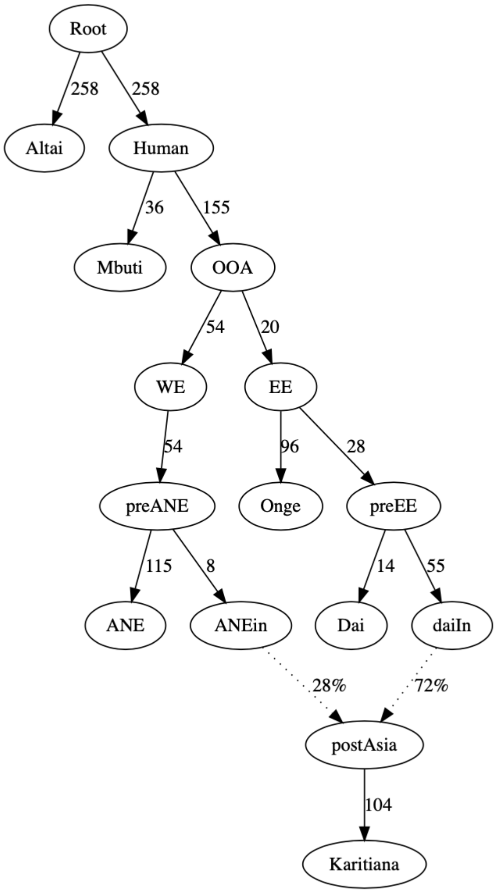


Figure 5: Drift lengths and admixture proportions inferred by `qpGraph` on the Eurasian-American dataset. The topology was pre-specified.

Similarly, the drift of 120 along the edge (2,5) corresponds to the OOA-WE and WE-pre-ANE drifts of 54 each ($54 + 54 = 108$).

At $K = 4$, we are unable to capture the full continuity of mixing proportions, and as a result, our drift lengths and admixture proportions do not match those of `qpGraph` exactly. Nonetheless, they are close: our inference of 25% and 75% are close to the precise values inferred by `qpGraph` of 28% and 72%, and the drift lengths match closely as well. The drift lengths around Karatiana are an exception, but is likely due to the fact that `qpGraph` cannot distinguish between drift that occurs before or after admixture.

Overall, `miqoGraph`'s ability to produce admixture graphs in seconds and match both `qpGraph`-enumerated results as well as knowledge from the literature [3] illustrates the power of mixed-integer quadratic optimization in fitting topologies, drifts, and admixture proportions jointly.

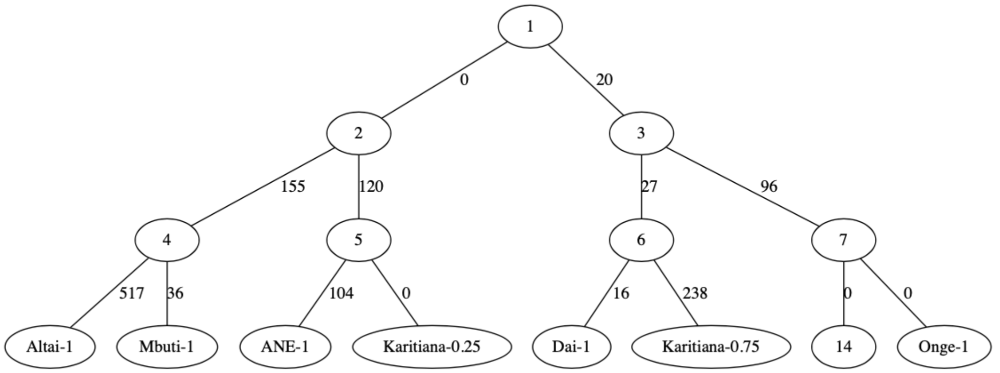


(a) Topology, drift lengths and admixture proportions inferred by `miqoGraph`. Nodes of the binary tree are labeled with number $1-15$, with the leaves corresponding to nodes $8-15$. The label "Altai-1" in node 8 means that Altai was assigned to node 8 at 100%; the label "Karitiana-0.25" in node 11 means that Karitiana was assigned to node 11 at 25%.

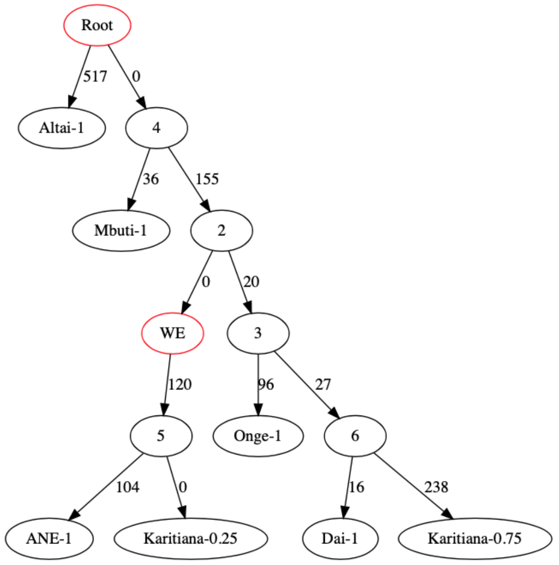


(b) An equivalent permutation of the binary tree in Figure 6a, with directedness added to the edges. Root and WE (red borders) were absorbed into the (4, Altai-1) and (2, 5) edges, respectively. Nodes 1, 7, and 14 do not appear because they are meaningless filler nodes.

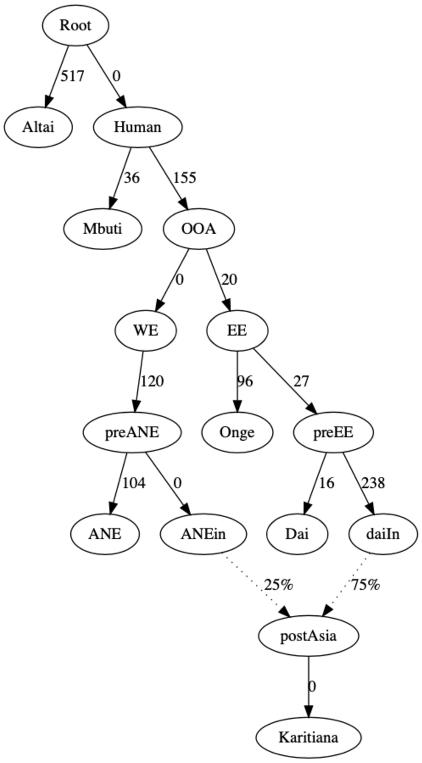


(c) The graph in Figure 6b with the ancestral nodes labeled according to Figure 5.

Figure 6: Topology, drift lengths and admixture proportions inferred by `miqoGraph` on the Eurasian-American dataset. Each subfigure represents a step in translating the output of `miqoGraph` to the format of `qpGraph`. Figure 6a shows the immediate binary-tree-formatted output produced by `miqoGraph`, which is permuted in Figure 6b, and then relabeled in Figure 6a. This process illustrates that the output from `miqoGraph` matches that produced by enumeration and `qpGraph`.
